# A tardigrade cytochrome P450 perturbs *Drosophila melanogaster* development

**DOI:** 10.1101/2025.05.31.657185

**Authors:** Caitlyn L Perry, Charles Robin

## Abstract

An analysis (Nelson, 2018) of *Hypsibius exemplaris* cytochrome P450 genes has revealed the existence of a gene (dubbed *Cyp18k1*) showing moderate sequence similarity with the ecdysteroid-inactivating arthropod Cyp18a group. Using a transgenic *H. exemplaris Cyp18k1* construct, we have demonstrated that ubiquitous CYP18K1 expression results in lethal failure of abdominal histoblast proliferation in *Drosophila melanogaster*.

## Description

Our understanding of early arthropod evolution is constrained by lack of information about arthropods’ closest living relatives, onychophorans and tardigrades (Giacomelli *et al*., 2025). While these lineages clearly show some innovations (e.g. loss of some Hox genes in tardigrades; Smith *et al*., 2016), when considered alongside more distant relatives, they can assist in clarifying the timing of events in arthropod and pre-arthropod evolution. This is particularly relevant to traits which are invariant within arthropods but absent from more distant ecdysozoan groups like nematodes, such as the role of ecdysteroids in regulating moulting (Chitwood and Feldlaufer, 1990).

Nelson’s (2018) analysis of the *Hypsibius exemplaris* cytochrome P450 (hereafter ‘P450’) inventory revealed a clear representative of the arthropod ecdysteroidogenic Cyp315 group, but no obvious one-to-one orthologs of the four other ecdysteroidogenic P450s. While the latter finding indicates that *H. exemplaris* is very unlikely to synthesise ecdysteroids, the former suggests that some form of sterol metabolism does occur in this species. Another *H. exemplaris* P450 – *Cyp18k1* – provides further evidence for such a pathway. This gene’s name reflects its moderate sequence-level similarity to arthropod Cyp18a genes, which encode ecdysteroid-26-hydroxylase enzymes catalysing the irreversible conversion of 20-hydroxyecdysone to hormonally inactive ecdysonoic acid (Bassett *et al*., 1997). Cyp18a genes are essential for viability in some arthropod species (e.g. *Drosophila melanogaster*, Guittard *et al*., 2011), but have been lost entirely from others (e.g. *Anopheles gambiae*; Feyereisen, 2006).

CYP18A enzymes form a sister clade to the ecdysteroidogenic CYP306A enzymes, which act as ecdysteroid-25-hydroxylases (Warren *et al*., 2004), and *Cyp18a1* and *Cyp306a1* are conserved in synteny across Mandibulata (Dermauw *et al*., 2020). Chelicerate genomes contain only single representatives of the combined CYP18A/CYP306A clade, which typically group with mandibulate *Cyp18a1* sequences in phylogenetic inference (e.g. Dermauw *et al*., 2020). However, the exceptionally rapid evolution of CYP306 genes observed in some arthropods (Good *et al*., 2014) may result in a misleading phylogenetic signal with respect to both the chelicerate and tardigrade Cyp18 genes. As we suspect that sequence similarity is not a reliable guide to *H. exemplaris* CYP18K1 function, we expressed *H. exemplaris Cyp18k1* in *D. melanogaster*, intending to determine its capacity to rescue mutations in arthropod ecdysteroid synthesis genes.

These efforts were frustrated by the fact that ubiquitous expression of *Cyp18k1* resulted in lethality even in the absence of mutations disrupting ecdysteroid synthesis; all (n = 277) adult offspring of an act-5C-GAL4/CyO-GFP x UAS-*Cyp18k1*/UAS-*Cyp18k1* cross were GFP-positive (indicating that they lacked the actin driver and thus did not express the transgene). We determined the lethal phase of *Cyp18k1* expression by identifying the stage at which no GFP-negative individuals could be found; this analysis revealed that expression of *Cyp18k1* had no obvious detrimental effect even at late pupal stages, but consistently prevented adult emergence, despite grossly normal development of structures such as eyes and wings within the pupal case. Removing the pupal case from GFP-negative pharate adults revealed specific failure of the abdominal epidermis to develop (see Figure 1). This was not the case when spookier-GAL4 (which drives expression in ecdysteroidogenic tissues; Moeller *et al*., 2017) was substituted for act-5C-GAL4, where a slight but not statistically significant excess of GFP-positive adults was observed (147 of 237, p = 0.12, two-tailed Fisher’s exact test). Driving *Cyp18k1* with the ubiquitous tubulin-GAL4 driver also resulted in only GFP-positive adults (n = 48). qPCR data did not indicate any difference in expression levels of the ecdysteroid-responsive genes E75 (p = 0.57, two-tailed t-test; see Supplemental Table 1) or E93 (p = 0.64, two-tailed t-test) between *Cyp18k1*-expressing flies and their GFP-positive siblings.

**Figure 1.**
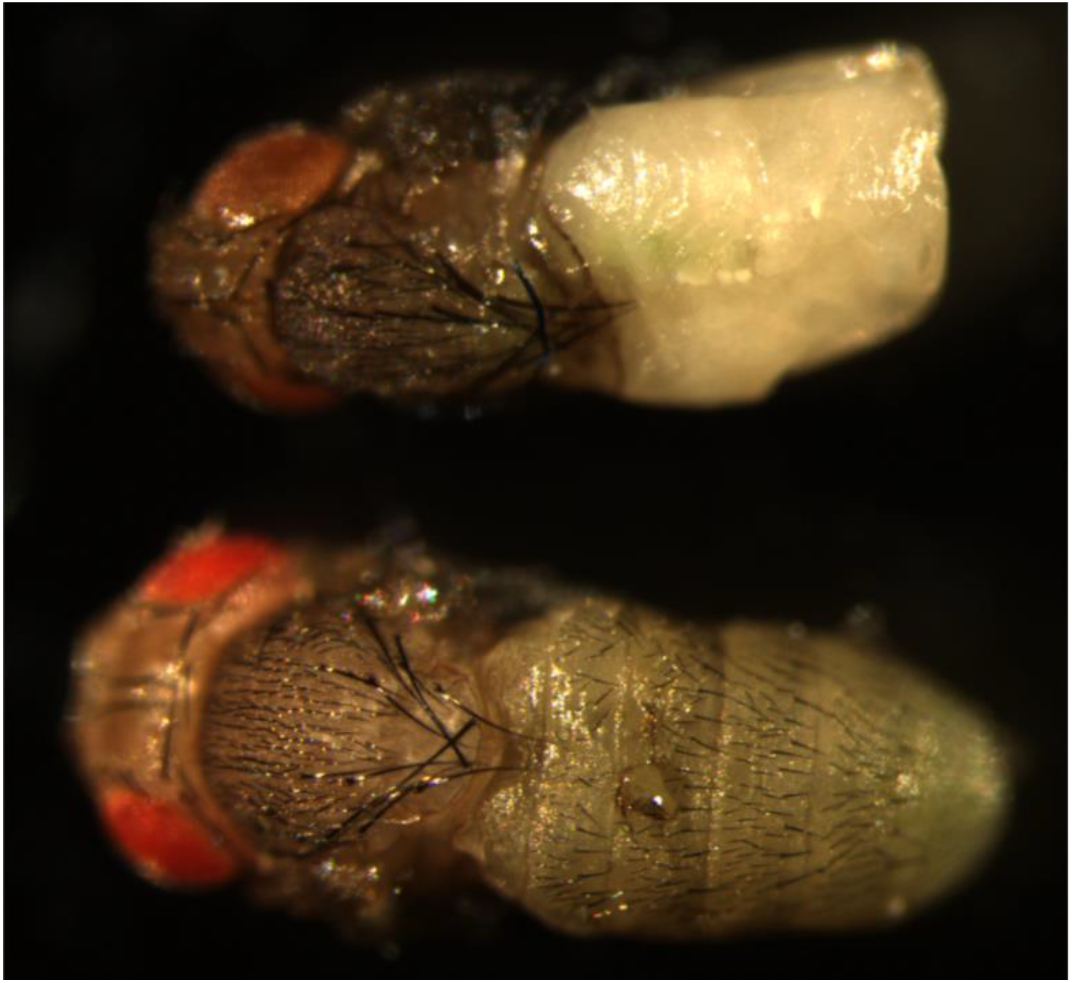
Pharate adults with the act-5C-GAL4/UAS-*Cyp18k1* (top) and CyO-GFP/UAS-*Cyp18k1* (bottom) genotypes. Normal eye and thorax development can be seen in the *Cyp18k1*-expressing fly, and the greenish meconium suggests typical breakdown of the larval gut.

The overexpression phenotype of *H. exemplaris Cyp18k1* differs notably from that of endogenous *D. melanogaster Cyp18a1* (Guittard *et al*., 2011). While ubiquitous *Cyp18a1* overexpression is also lethal (Guittard *et al*., 2011), in this case death occurs at late embryonic stages, presumably due to the role of ecdysteroids in embryogenesis (e.g. Bownes *et al*., 1988). Neither the specificity of the *Cyp18k1* overexpression phenotype nor the normal expression of ecdysteroid-induced genes in *Cyp18k1*-positive larvae supports globally reduced ecdysteroid levels as the cause of lethality. However, the *D. melanogaster Cyp18k1* overexpression phenotype resembles the effects of abdominal histoblast proliferation failure (Sekyrova *et al*., 2010), and abdominal histoblast development is known to be ecdysteroid-dependent (Ninov *et al*., 2007).

These observations might be reconciled by the hypothesis that CYP18K1 inactivates ecdysteroid derivatives with a specific role in *D. melanogaster* abdominal histoblast proliferation. This is not an entirely *ad hoc* suggestion, as the abdomen is the only adult structure which does not begin proliferation until late larval stages (Mandaravally Madhavan and Madhavan, 1980; Truman and Riddiford, 2002). Given evidence that 3-dehydroecdysteroids are synthesised in third-instar *D. melanogaster* larvae (Sommé-Martin *et al*., 1988), these are an (admittedly speculative) candidate for a specific role in abdominal histoblast development.

Several developmental transcriptomic series from *H. exemplaris* (Levin *et al*., 2016; Yoshida *et al*., 2017) indicate substantial fluctuations in CYP18K1 expression level during egg development, consistent with an endogenous developmental function, but our near-total ignorance of tardigrade sterol biology is a barrier to even proposing (let alone testing) candidate roles for this enzyme. Identification of the molecules on which CYP18K1 acts in *D. melanogaster* would be a useful first step towards clarifying a possible role for steroid-derived molecules in tardigrade development.

## Methods

Whole-plasmid synthesis of *Hysibius exemplaris Cyp18k1* in a pUAST-attB backbone was undertaken by VectorBuilder, and injections of this plasmid into a *D. melanogaster* line expressing ϕ31 integrase were performed by BestGene Inc. Expression was driven by act-5C-GAL4 (Ito *et al*., 1997), tubulin-GAL4 (Lee and Luo, 1999) or spookier-GAL4 (Moeller *et al*., 2017). *D. melanogaster* stocks were obtained from the Bloomington *Drosophila* Stock Center. All stocks were reared at 25°C on molasses media.

In order to determine how expression of *H. exemplaris Cyp18k1* affects ecdysteroid signalling, expression levels of the ecdysteroid-responsive *E75* (Segraves and Hogness, 1990) and *E93* (Baehrecke and Thummel, 1995) were measured, along with control genes showing a peak of expression at a similar time but not known to be ecdysteroid-regulated (*chaoptin, icarus* and *papillote*, identified using FlyBase RNA-Seq Query Tools). RNA was isolated from nine act-5C-GAL4/UAS-*Cyp18k1* and nine CyO-GFP/UAS-*Cyp18k1* four-day-old pupae (genotype inferred from fluorescence) by chloroform extraction, reverse-transcribed and used in quantitative PCR reactions (normalised against *rpl11*).

## Reagents

### Oligo sequences

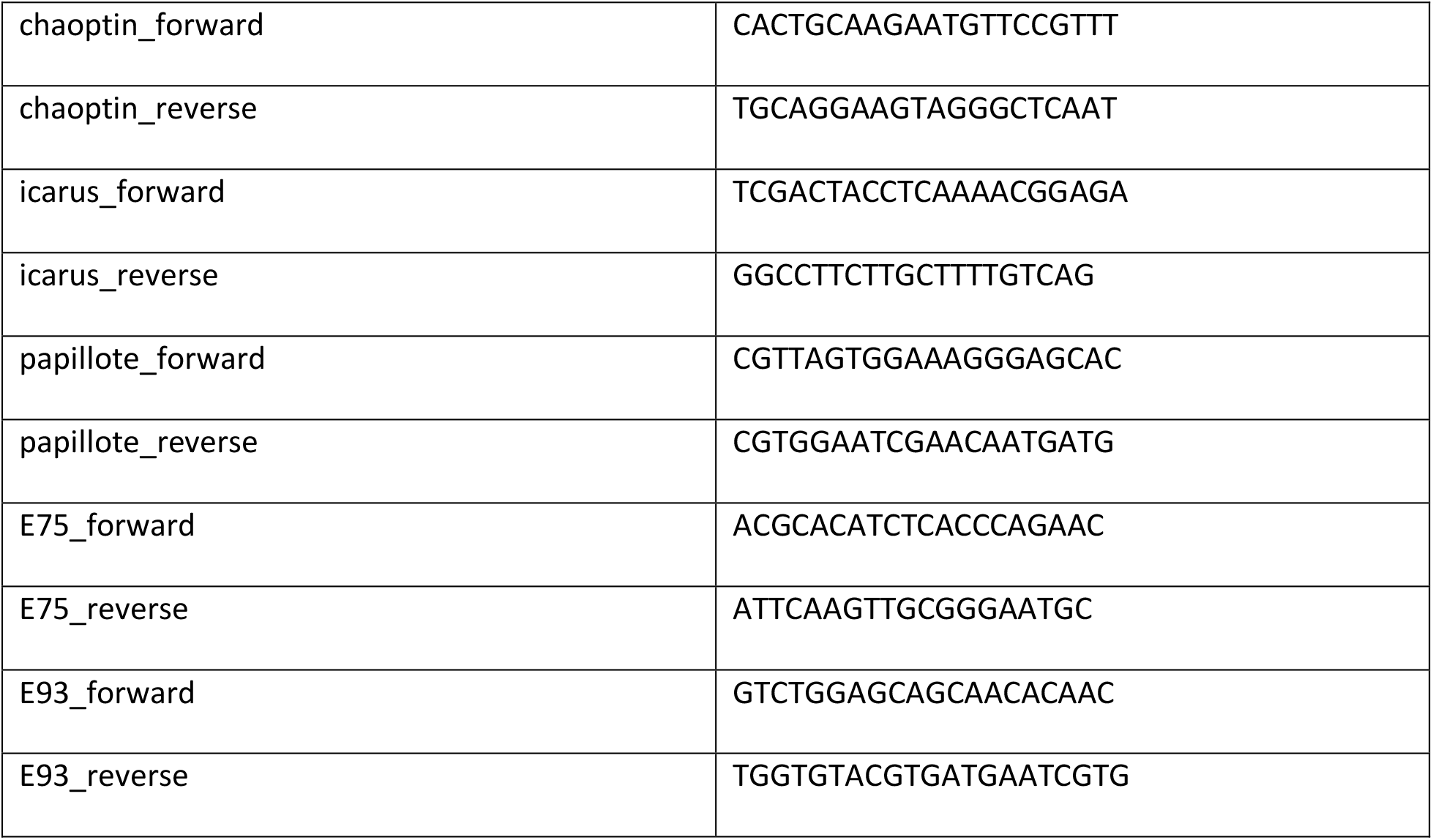

### Fly lines

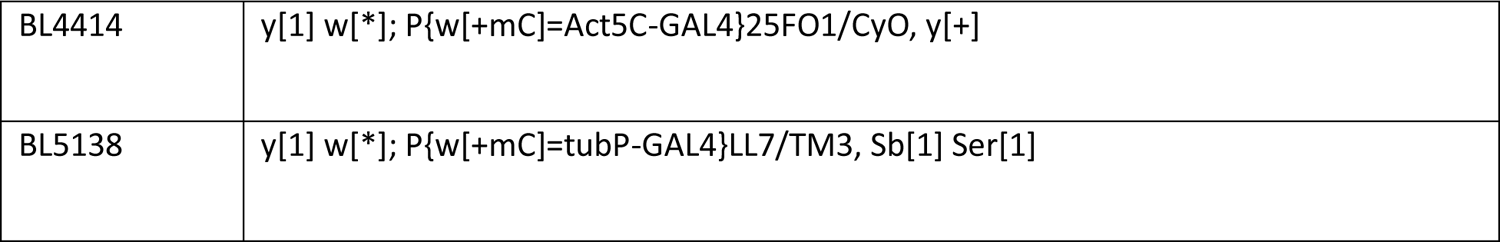
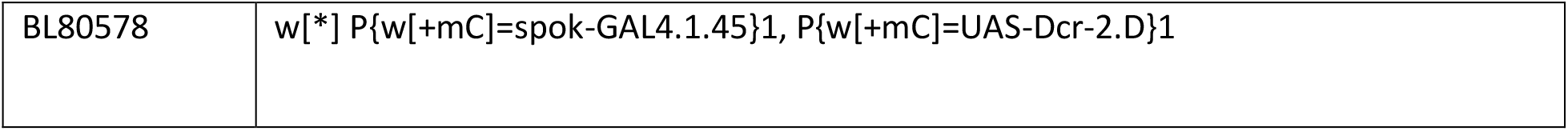

Fly lines:
|  |  |
| --- | --- |
| BL4414 | y[1] w[*]; P{w[+mC]=Act5C-GAL4}25FO1/CyO, y[+] |
| BL5138 | y[1] w[*]; P{w[+mC]=tubP-GAL4}LL7/TM3, Sb[1] Ser[1] |
| BL80578 | w[*] P{w[+mC]=spok-GAL4.1.45}1, P{w[+mC]=UAS-Dcr-2.D}1 |

### qPCR reagents

TRIsure (Bioline), DNase I (Thermo Fisher Scientific), GoScript Reverse Transcriptase kit (Promega), SsoAdvanced Universal SYBR Green Supermix (Bio–Rad), CFX96 C1000 Touch System (Bio–Rad).

